# A robust sequencing assay of a thousand amplicons for the high-throughput population monitoring of Alpine ibex immunogenetics

**DOI:** 10.1101/2020.10.27.357194

**Authors:** Camille Kessler, Alice Brambilla, Dominique Waldvogel, Glauco Camenisch, Iris Biebach, Deborah M Leigh, Christine Grossen, Daniel Croll

## Abstract

Genetic variation is a major factor determining susceptibility to diseases. Polymorphism at the major histocompatibility complex (MHC) and other immune function loci can underlie health and reproductive success of individuals. Endangered species of low population size could be severely compromised to evolve disease resistance due to reduced adaptive variation. A major impediment to screen adaptive genetic variation in wild species is the difficulty to comprehensively genotype immune-related loci based on low input material. Here, we design and validate a targeted amplicon sequencing assay to parallelize the analysis of a thousand loci of the MHC, other immunity-related genes, and genome-wide markers for the assessment of population structure. We apply the approach to Alpine ibex, one of the most successful examples of restoration of a large mammal in Europe. We used 51 whole genome sequenced individuals to select representative target SNPs. We integrated SNP call data from four related species for amplification robustness and genotyped 158 Alpine ibex individuals for validation. We show that the genome-wide markers perform equally well at resolving population structure as RAD-seq or low-coverage genome sequencing datasets with orders of magnitude more markers. The targeted amplicon sequencing assay is robust to >100-fold variation in input DNA quantity and generates useful genotype information from fecal samples. The amplicon marker set also identified recent species hybridization events with domestic goats. The immune loci show unexpectedly high degrees of differentiation within the species. Our assay strategy can realistically be implemented into population genetic surveys of a large range of species.

## Introduction

Biodiversity is currently undergoing a dramatic decline caused by ecological and anthropological pressures (Barnosky et al., 2011; G. Ceballos et al., 2015; Gerardo Ceballos et al., 2017; Wwf, 2018). Endangered species are particularly prone to genetic risks due to population bottlenecks and habitat fragmentation that lead to low genetic diversity, inbreeding, introgression from related species and deleterious mutation accumulation (Allendorf et al., 2010; Frankham, 2005). Low genetic diversity can affect fitness and survival, as shown for instance in cheetah, Florida panthers and Alpine ibex (Brambilla et al., 2015; O’Brien et al., 1983, 1985; Pimm et al., 2006; Reed & Frankham, 2003; Roelke et al., 1993). Loss of genetic diversity at adaptive immune loci is particularly problematic because populations will lack genetic variants conferring disease resistance, in particular infectious diseases, as it has been shown for example in amphibians (Kosch et al., 2019; Savage et al., 2011), Tasmanian devils (Siddle et al., 2007), giant panda (Zhu et al., 2020) and Alpine ibex (Brambilla et al., 2018). However, despite the importance for conservation management, the genetic underpinnings of disease susceptibility are largely unknown in most non-model species (Schoville et al., 2012). The major obstacles in investigating wildlife-immunogenetics are the challenge of collecting quantitatively and qualitatively adequate phenotypic data on diseases in wild populations, as well as the lack of genetic tools suitable in the conservation framework (Holderegger et al., 2019). Hence, there is a pressing need to establish effective genetic monitoring tools for many wild species (Acevedo-Whitehouse & Cunningham, 2006; Allendorf et al., 2010).

The major histocompatibility complex (MHC), a highly polymorphic region whose products are involved in foreign antigen recognition, is an important player for disease susceptibility in vertebrates. Co-evolution with pathogens often causes balancing selection on MHC polymorphism, maintaining high genetic diversity including deeply divergent alleles. Genetic variants under selection often encode the ability to present a wider range of antigens to T-cells and, thus, to recognize a greater variety of parasites. Consequently, heterozygotes often have elevated resistance and are favored by selection (Bernatchez & Landry, 2003). Species that underwent a strong bottleneck such as Alpine and Iberian ibex (*Capra ibex and C. pyrenica*), Tasmanian devils, Cheetahs or Galapagos penguins (*Spheniscus mendiculus*), show strongly reduced genetic diversity at the MHC compared to related species (Angelone et al., 2018; Bollmer et al., 2007; Brambilla et al., 2018). Low genetic diversity in cheetahs is thought to have contributed to high mortality rates (O’Brien et al., 1983; 2017). High prevalence of a fatal facial cancer in Tasmanian devil is thought to, at least, partially stem from low MHC diversity (Siddle et al., 2007). Low levels of MHC diversity are consequently a major threat for endangered species. Alongside with the adaptive immune system, the innate immune system plays an important role in the defense against a wide range of pathogens. Genetic variation within some genes, *e.g.* Toll-like receptors (TLR), has been shown to be linked with variation in immune competence (Ammerdorffer et al., 2014; Tschirren et al., 2013). Yet the field of wildlife-immunogenetics is only emerging and immune-related genes outside of the MHC remain heavily understudied in wild species. The major challenge for monitoring non-model wild species is that inferring causal disease susceptibility from other species is only rarely possible and associations need to be established for each species – pathogen interaction (Acevedo-Whitehouse & Cunningham, 2006).

To make informed management decisions, the genetic health of populations should also be taken in account and assessed (Allendorf et al., 2010). Integrating information about immune-related polymorphisms and genetic diversity across the genome remains a major challenge. Conservation genetics studies largely rely on microsatellites (Allendorf et al., 2010; Brambilla et al., 2015; Ouborg et al., 2010; Witzenberger & Hochkirch, 2014), which are inexpensive and applicable for a wide range of input material. However, only in exceptional cases do microsatellites reveal adaptive genetic variation and are thus not applicable to most relevant immune-related polymorphism (Allendorf et al., 2010; Steiner et al., 2013). Similarly, modern conservation genomics methods based on restriction site associated DNA sequencing (e.g. RAD, ddRAD, GBS, Davey et al., 2011) intrinsically lack the power to target particular polymorphisms of interest in the genome (Garvin et al., 2010). Whole genome sequencing solves the issue of marker selection but is expensive and requires high-quality input material to work reliably (Davey et al., 2011). Conservation genetics monitoring requires targeted genotyping assays that tolerate a wide range of input material, assess specific polymorphisms of interest and is cost effective.

Here, we develop and validate a high-throughput amplicon sequencing array, which enables immunogenetic monitoring of Alpine ibex. The species is an excellent model to assess the broad applicability of novel genotyping assays for conservation because of a dramatic recent bottleneck caused by a near extinction event in the 19th century (Grodinsky & Stüwe, 1987). The census size was reduced to less than 100 individuals limited to one single population in Northern Italy (Gran Paradiso). A captive breeding program restored the species to a current census of 53'000 across the European Alps (Brambilla et al., 2020). The historic bottleneck left substantial genome-wide signatures of low heterozygosity, in particular at MHC loci, which may threaten long-term population viability (Grossen et al., 2014, 2018, 2020). Hybridization events with domestic goat produced introgression tracts at the MHC re-establishing some genetic variation, which may have been lost due to the species bottleneck (Grossen et al., 2014). Recently, concerns were raised over population declines as direct and indirect consequences of epizootic disease outbreaks (*e.g.* sarcoptic mange, respiratory diseases, infectious keratoconjunctivitis, brucellosis). One recent disease outbreak concerned cases of brucellosis in the area of Bargy (France). Additional cases were identified in bovine and humans and were considered a broader threat (Mick et al., 2014). As the ibex were seen as a reservoir for the brucellosis pathogen, French authorities undertook a massive eradication program leading to the culling of more than 250 individuals (44% of the estimated population) within two years following the outbreak (Mick et al., 2014; Quéméré et al., 2020). Hence, Alpine ibex present a model species where high-throughput monitoring of inbreeding levels, potential hybridization events and immunogenetic diversity can substantially improve genetic analyses and conservation management efforts.

We established a high-throughput assay of nearly a thousand loci covering largely genome-wide (putatively neutral) polymorphisms, variants found at the MHC and other immune-related loci as well as diagnostic variants useful to detect recent hybridization events with domestic goat. We used 51 whole-genome sequences of Alpine ibex and domestic goat to identify relevant SNPs to be targeted for the amplicons. Furthermore, we used whole genome sequences to mask polymorphism at primer sites to maximize amplification success. Based on highly parallel Illumina amplicon sequencing of 172 Alpine ibex, Iberian ibex and domestic goat samples, we assessed the accuracy and robustness of the assay across highly variable input sample quality. Finally, we compared the high-throughput assay with RAD-seq and low-coverage whole-genome sequencing datasets on the same populations.

## Materials and methods

### Collection of individuals and DNA extraction

Genotyping was performed on 158 Alpine ibex samples, representing 8 populations with 19-20 individuals each. The samples included also an individual suspected to be an albino. Sample material consisted either of tissue or blood collected either by biopsy darting, during captures or legal hunting (Biebach & Keller, 2009; Willisch et al., 2012). We included four fecal samples collected in the Gran Paradiso National Park as well as six individuals representing suspected hybrids based on field observations, five domestic goats (*Capra hircus*) and three Iberian ibex individuals. Detailed information about the origin, collection method and sampling year of each sample is provided in Supplementary Table S1. All DNA extractions were carried out using the DNeasy Blood & Tissue kit (QIAGEN).

### SNP discovery based on whole-genome sequenced individuals across species

We used 29 Alpine ibex whole genome sequences (representing seven different populations) to identify segregating SNPs to design our target amplicons. 36 additional genome sequences (4 Iberian ibex, 16 domestic goats, 6 bezoar, 2 Siberian ibex, 2 Nubian ibex, 1 Markhor and 5 sheep, see (Grossen et al., 2020) for details) were used to detect diagnostic markers and to mask highly polymorphic regions (see below). Data from domestic goat, bezoar and sheep were produced by the NextSeq Consortium. Trimmed reads (Trimmomatic v.0.36, (Bolger et al., 2014) were mapped using bwa-mem (Li et al., 2009) to the domestic goat reference genome (version ARS1, Bickhart et al., 2017) and duplicates were marked using MarkDuplicates from Picard (http://broadinstitute.github.io/picard, v.1.130). After genotype calling using HaplotypeCaller and GenotypeGVCF (McKenna et al., 2010; GATK, v.4.0.8.0, Van der Auwera et al., 2013), SNPs were removed using VariantFiltration of GATK if: QD <2.0, FS > 40.0, SOR > 5.0, MQ < 20.0, −3.0 > MQRandkSum > 3.0, −3.0 > ReadPosRankSum > 3.0 and AN < 46 (80% of all Alpine ibex individuals). We identified a total of 138 million SNPs segregating among all analyzed species with 5.3 million SNPs segregating among the 29 sequenced Alpine ibex. Genome-wide polymorphism was used in three ways to design amplicons for the high-throughput sequencing assay: (1) To discover SNPs variable in Alpine ibex to be targeted for amplification, (2) to discover SNPs differentially fixed between Alpine ibex and domestic goats (diagnostic markers). (3) to mask polymorphic sites near targeted SNPs to prevent designing primers on polymorphic sites potentially causing amplification drop-outs.

### SNP effect prediction and selection

We annotated SNPs using SnpEff v4.3t 2017-11-24 (Cingolani et al., 2012) with gene annotations produced for the domestic goat genome ARS1 as reference (Bickhart et al., 2017). SNPs located in repeat-masked regions of the reference genome were excluded to avoid designing amplicons in repetitive regions. We designed four types of marker sets.

Marker type 1 - Genome-wide, putatively neutral markers: For the design of genome-wide markers, we focused on SNPs segregating only among Alpine ibex individuals from the Gran Paradiso population, the source of all current Alpine ibex populations (4 million SNPs). Our aim was to most accurately reflect segregating neutral polymorphism in the species. Hence, including non-source individuals could lead to ascertainment bias. Loci were selected with a genotyping rate of ≥70%, a minimal genotyping quality of 20 and at a minimal distance of 2.3 Mb between adjacent SNPs.

Marker type 2 - To detect recent hybridization events, we identified loci with fixed alleles distinguishing domestic goats and Alpine ibex. Specifically, we chose loci with fixed allele frequency differences between all Alpine ibex and all goats (both domestic goat and bezoar), with a minimal genotyping rate of 80%, a minimal genotyping quality of 20 and a minimal distance between adjacent SNPs of 13 Mb. Markers in genes relevant to the immune system were selected for two subgroups (here called MHC region and Immune, outside MHC) with a minimal allele count of 1 among Alpine ibex.

Marker type 3 in MHC region: Because of the importance of the MHC region for immune functions and evidence for introgression from the domestic goat, we covered the entire MHC region on chromosome 23 (positions 20,892,916-23,588,623 bp). The chromosomal location of the MHC was identified using BLASTN v2.7.1+ (Altschul et al., 1990) based on gene sequences reported as belonging to the MHC in a previous goat genome assembly version (CHIR1, Dong et al., 2013, Supplementary Table S17). Additionally, we used gene ontology (GO) and gene homology to search for all MHC-related genes on chromosome 23. Matching gene sequences within 250 kb of the homologous MHC region of the CHIR1 assembly were considered as part of the MHC region for further analyses. We split the MHC region into 270 windows of 10 kb using bedtools v2.27.1 (Quinlan & Hall, 2010) to identify SNPs for the amplicon assay (Figure 1B). Evidence for introgression from domestic goat is particularly strong at the *DRB* exon 2 (Grossen et al., 2014). We hence designed the amplicon assay with more dense SNPs in the *DRB* region (positions 23,411,211-23,511,211, Figure 1B). The *DRB* was localized between positions 23,451,944-23,470,477 using BLASTN v2.7.1+ (Altschul et al., 1990) and the *DRB* sequence provided by (Dong et al., 2013). A buffer zone of 40'733 bp before and after the gene was added thereby also including gene ENSCHIG00000008942, which encodes immune-related functions (Figure 1B, lower panel). We divided the *DRB* region into 20 windows of 5 kb. We randomly selected a single SNP from every window in the defined MHC and *DRB* by prioritizing MODERATE impact mutations based on Snpeff and SNPs in coding regions. We manually selected a marker in the *DRB* exon 2 at position 23,460,796 bp.

**Figure 1:**
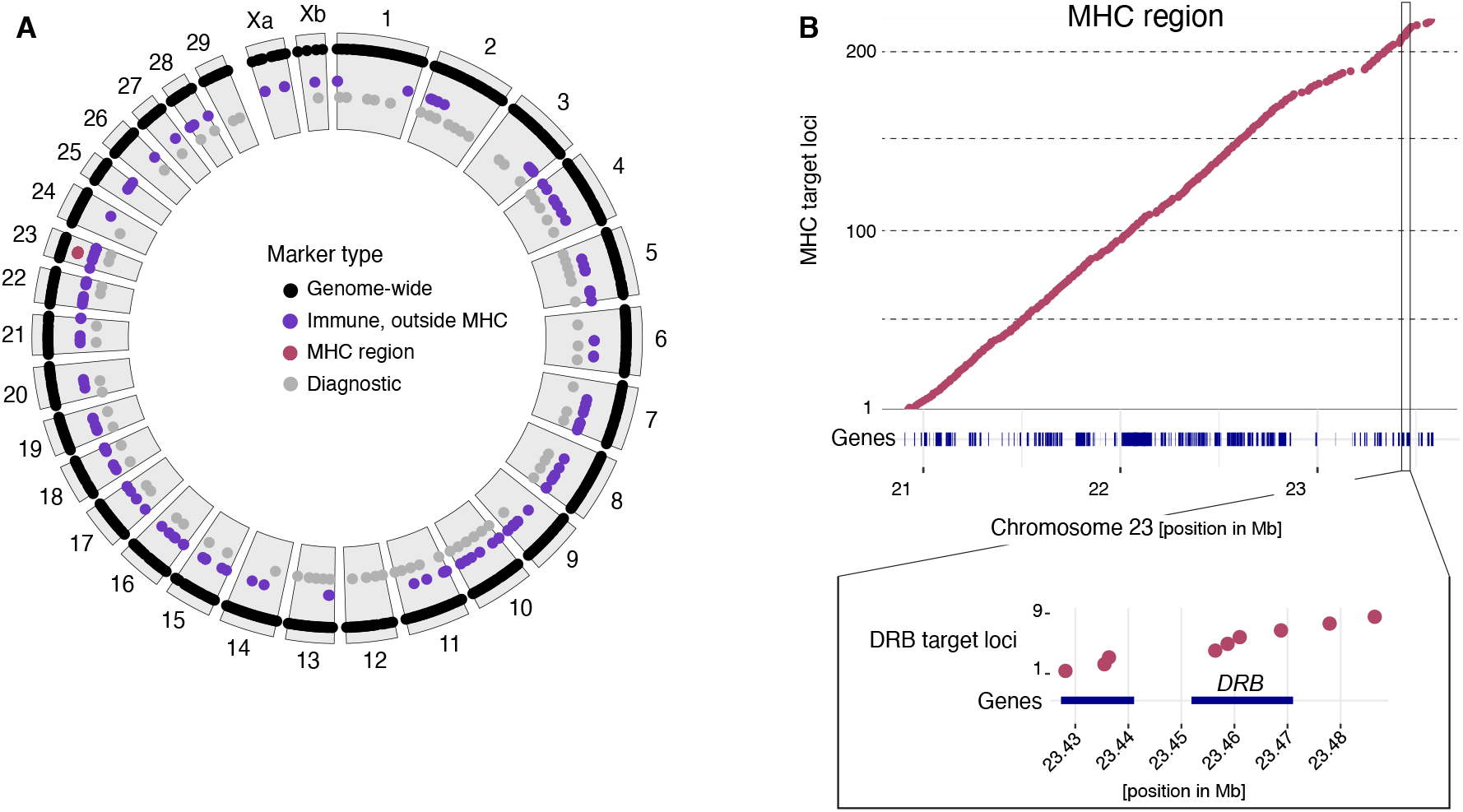
Design and distribution of targeted amplicon sequencing loci. A) Distribution along chromosomes for each marker category (genome-wide, immune loci, MHC region, diagnostic); B) Regularly spaced loci across the MHC region on chromosome 23 (20.89-23.59 Mb). Blue segments identify coding sequences in the MHC. The *DRB* gene region of the MHC (zoom view) was targeted by a denser array of loci.

Marker type 4: We targeted immune-related genes, outside of the MHC region on chromosome 23, based on a candidate gene approach using gene ontology (GO), gene homology, as well as literature reports (Acevedo-Whitehouse & Cunningham, 2006; Turner et al., 2011). We searched for all MHC-related GO terms on Ensembl Biomart in the goat genome. Then, we performed InterProScan v. 5.31-70.0 analyses and searched for protein domain annotations matching the term “histocompatibility”. For the lists by Acevedo-Whitehouse and Cunningham (2006) and Turner et al. (2011), we focused on 211 genes encoding important immune functions for mammals including interferons, interleukins, toll-like receptors (TLR) and MHC-related genes outside of the MHC region on chromosome 23. Each retained gene locus was targeted by a single SNP prioritizing SNPs in coding sequences or with a MODERATE impact annotation based on SnpEff (Supplementary Table S2).

### Amplicon design

For each amplicon to be designed, we extracted a 1001 bp sequence from the domestic goat reference genome centered around the target SNP (IUPAC encoded) using bcftools v1.9 (https://www.htslib.org/). The extracted sequences had masked positions for all repetitive regions (masked reference genome) and positions which were polymorphic in Alpine ibex and/or with a minor allele count of three among all other *Capra* species (samtools v. 1.9 with option −m; (Li et al., 2009)). Sequences with more than 50% masked bases or sequences overlapping between different amplicon sets were excluded. We obtained a set of 1589 sequences for assay primer design by Fluidigm Inc. About 25% of the sequences were rejected by Fluidigm due to the lack of primer options matching the design criteria, leaving 1265 sequences for oligonucleotide primer synthesis (Figure 1). The targeted amplicon length was 200 bp.

### Targeted DNA sequencing library preparation and SNP calling

We prepared libraries following the manufacturer protocol “Library Preparation with the LP 192.24 IFC” using the Fluidigm Inc. Juno system, except for the minimal concentration of genomic DNA. Due to very low reaction volumes, the manufacturer recommends ≥ 100-200ng/μl. As most samples wouldn’t have reached that, we decided to use a minimal concentration of 50ng/μl. Samples also failing the concentration of 50ng/μl of genomic DNA, were concentrated using Ampure magnetic beads, whereas those, for which concentration to 50ng/μl was not possible due to low starting volumes, were used at the original concentration.

Barcoded and quality checked libraries were sequenced on a single lane of an Illumina NextSeq 500 in mid-output mode adding ~30% PhiX to avoid potential problems due to low sequence complexity. We demultiplexed raw read data using bcl2fastq v2.19.0.316 and used Trimmomatic v0.38 (Bolger et al., 2014) for quality trimming. Forward and reverse reads were merged using FLASH v1.2.11 (Magoč & Salzberg, 2011) and aligned to the goat reference genome ARS1 using bowtie2 v2.3.5 (Langmead & Salzberg, 2012). Read depths for each step were estimated with MultiQC v.1.7 (Ewels et al., 2016). We called SNPs using HaplotypeCaller, CombineGVCFs and GenotypeGVCFs from GATK v4.0.1 (McKenna et al., 2010; Van der Auwera et al., 2013). Variant sites were further filtered to meet the following conditions: QD < 5, MQ < 20, −2 > ReadPosRankSum > 2, −2 > MQRankSum > 2, −2 > BaseQRankSum > 2.

### Marker system performance on Alpine ibex populations

To evaluate key performance metrics of the new high-throughput amplicon sequencing assay, we compared the outcome against two major classes of current population genomics sequencing approaches: RAD-seq and low-coverage whole genome sequencing. We used datasets reporting analyses on four populations also included in our study (Grossen et al., 2018, 2020; Leigh et al., 2018): the founder population of Gran Paradiso and the three Swiss populations Pleureur, Brienzer Rothorn and Albris. Because high-coverage whole genome sequencing is not feasible for large scale, practical applications, we generated realistic, low-coverage datasets at approximately 1x coverage. For this, we downsampled 15 whole genome Illumina sequencing datasets (Grossen et al., 2020). We used sambamba v.0.6.6 (Tarasov et al., 2015) to downsample individual bamfiles to a fraction of 0.05 (producing a final coverage of ~1x). We used the software ANGSD (Korneliussen et al., 2014) to calculate genotype likelihoods with the following options: -doGlf 2, -doMajorMinor 1, -doMaf 1, - minMaf 0.05, -SNP_pval 1e-6, -minMapQ 20, -minQ 20, -skipTriallelic 1, -uniqueOnly 1, -remove_bads 1, -only_proper_pairs 1. The resulting likelihoods were used to run PCAngsd (Meisner & Albrechtsen, n.d.) and NGSadmix (Skotte et al., 2013). For the RAD-seq dataset, we first trimmed reads using Trimmomatic v3.6 (Bolger et al., 2014) and performed read mapping using Hisat2 v.2.1 (Kim et al., 2019) on the ARS1 reference genome. We de-duplicated bam files using Markduplicates from Picard v. v.2.5 (http://broadinstitute.github.io/picard/) and called SNPs on all autosomes (1-29) using the GATK v 4.1 pipeline with HaplotypeCaller, GenomicsDBimport and GenotypeGVCFs (McKenna et al., 2010). SNPs were flagged using the GATK VariantFiltration tool if any of the conditions were matched: QD <2.0, FS > 60.0, SOR > 3.0, MQ < 30.0, −12.5 > MQRandkSum > 12.5 and −8.0 > ReadPosRankSum. Next, we filtered for SNPs falling within 100 bp of a *SbfI* restriction cut site identified by *in silico* analyses with the ENSEMBL tool *restrict* (Yates et al., 2019). To compare genotyping performance between RAD-seq and the amplicon sequencing datasets, we filtered for a minimal individual genotyping rate of 0.5 and kept polymorphic sites only. We retained for the amplicon sequencing 892 SNPs genotyped in 75 individuals and for the RAD-seq 26,547 SNPs genotyped in 82 individuals. For all further analysis (population differentiation), we kept of the RAD seq dataset only bi-allelic SNPs with a minimal genotyping rate of 0.9, a minor allele frequency of 0.01 and a heterozygosity below 0.8 (*n =* 8316 SNPs retained). For the amplicon sequencing, we only retained SNPs from the genome-wide (neutral) set and removed sites on the Xa/b sex chromosome (*n* = 588 SNPs).

### Genetic data analyses

Genetic data analyses were done using R 4.0.2 (R Core Team 2018). The R package {BioCircos} was used to generate the circular plot. Principal component analyses (PCAs) were performed using the glPCA function from the R package {adegenet}. The R package {hierfstat} was used for FST calculations. The SNP intersection matrix was visualized with {UpSetR}. Genotype assignment plots were generated using sparse non-negative matrix factorization algorithms as implemented in the R package {LEA}. For each marker set, we ran 100 repetitions per K (K = 1-10) with entropy=TRUE to find the most likely number of clusters (*i.e.* K with the lowest entropy). Tajima’s *D* estimates in coding sequences (all immune-related genes represented on the amplicon) were calculated using the 29 Alpine ibex whole-genome sequencing datasets and the R package {PopGenome}.

## Results

### Assessment of locus quality across the targeted sequencing assay

We performed targeted amplicon sequencing of 1265 SNP loci covering genome-wide polymorphism and variants related to immune functions. The Fluidigm Inc. Juno microfluidics systems enables highly parallel amplification of all loci. We analyzed a total of 187 samples (representing 172 individuals) in a single run including 158 Alpine ibex, 5 domestic goats, 3 Iberian ibex and 6 suspected hybrids between Alpine ibex and domestic goats (Supplementary Table S1). HiSeq Illumina sequencing generated a total of 108 million read pairs after removal of PhiX spike-ins (32.4 Gbp total data). To assess genotyping quality across loci, we first focused only on samples that satisfied the manufacturer’s recommended DNA concentration of ≥50 ng/μl. Furthermore, we required that each analyzed individual produced at least 100’000 mapped reads across all loci (Figure 2A). The 65 retained individuals represented 7 different Alpine ibex populations, 2 domestic goat breeds as well as 3 Iberian ibex. The number of mapped reads ranged from 230’801-1’576’615 reads (Supplementary Table S3). Based on these high-quality samples, we found that the median read depth per locus was high with nearly all loci having >20 reads (median across loci = 389 reads). We found 7 loci with a median read depth of 0. The highest median read depth was 1931 for a marker designed in the MHC. Next, we analyzed the genotyping rate across loci and found that 940 loci were genotyped in all high-quality samples and 989 loci were genotyped in more than 75% of samples (Figure 2C). We discarded 28 loci with a genotyping rate below 75% to prioritize loci providing the highest information content across individuals. We retained a total of 989 high-quality loci for further analysis (Figure 2D).

**Figure 2:**
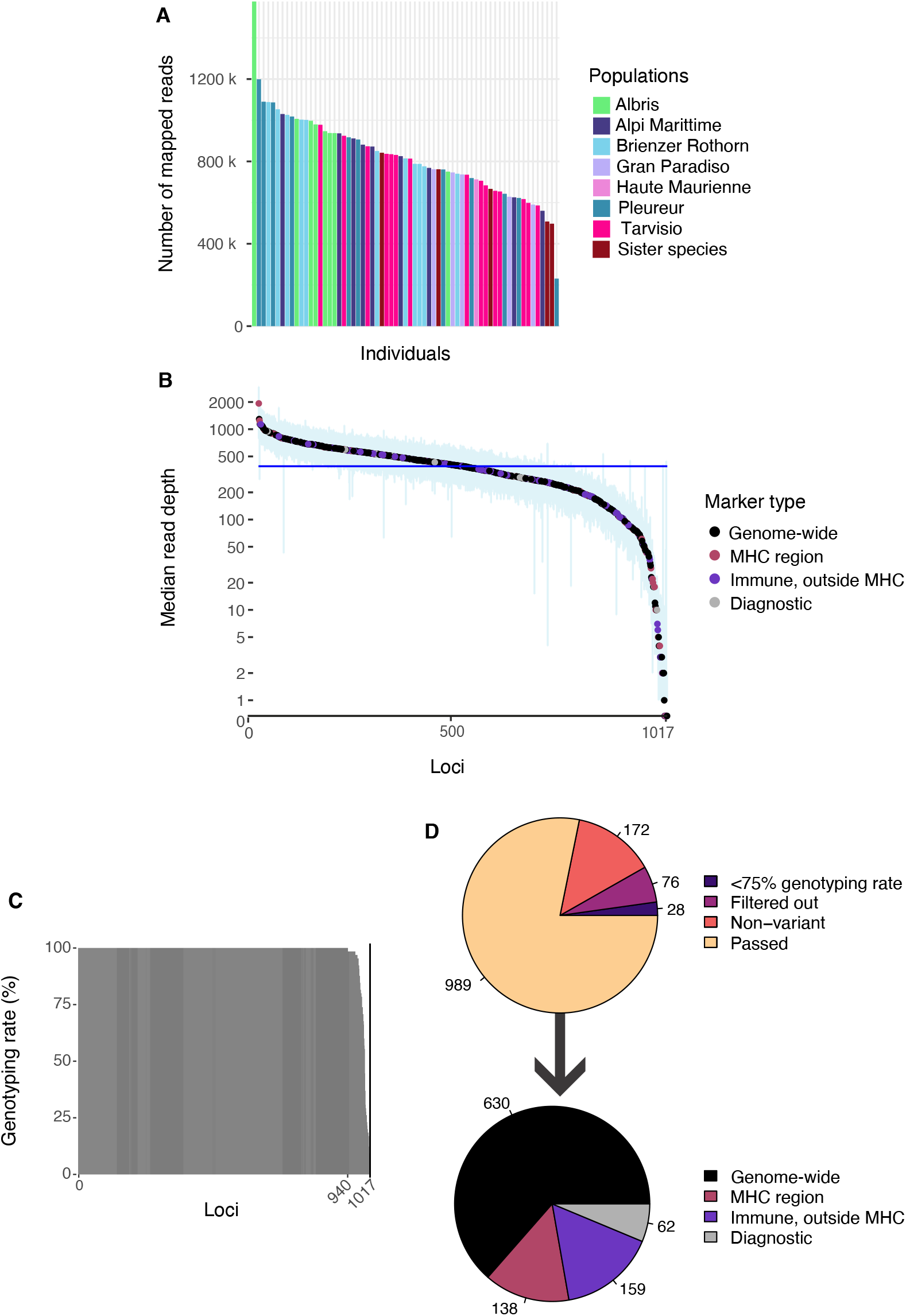
Assessment of amplification consistency and genotyping rates. A) Number of mapped reads per individual sample, colored by population *(N*=*65)*. Seven Alpine ibex populations are shown in addition to domestic goats (*N*=*2*) and Iberian ibex (*N*=*3*). Low-quality input DNA samples (fecal samples, etc.) were excluded. B) Median read depth per locus across 1017 amplicon loci. Markers are colored according to their category. The blue line represents the median (389) and the blue area represents the 95% confidence interval for each marker. C) Variation in genotyping rates across the 1017 loci. D) Outcomes of the different filtering stages. A total of 989 loci were retained for further analyses. The marker category of the retained markers is shown below.

### Assessment of genotyping rate and accuracy

The mean genotyping quality (GQ) across the 65 high-quality samples was on average 96 (Figure 3A). Taking advantage of four Alpine ibex individuals, which were both whole-genome sequenced and genotyped using our assay, we analyzed the overall accuracy of genotypes. Among the four individuals, a total of 62 loci were not assigned a genotype based on whole-genome sequencing. On average, 901 loci (range: 883-910, Figure 3B) showed perfectly matching genotypes and twenty loci (range: 17 - 27, Figure 3B) showed one mismatching allele (*i.e.* heterozygote vs. homozygote mismatch). The average genotype quality of the mismatched genotypes was 63 while it was 96 for matching genotypes. We found no complete allelic mismatch in any of the four individuals (*i.e.* homozygous calls for distinct alleles). An effective high-throughput genotyping assay should perform sufficiently well for low sample input quantity and quality. To establish a benchmark for input DNA sensitivity, we analyzed dilution series of input DNA. We found that genotyping rates >90% across loci are retained by diluting samples 25-fold from 100 ng/μl down to 4 ng/μl (Figure 3C). At 0.8 ng/μl, the genotyping rate was >75% for two out of three samples. The genotyping rate was >23% for 0.16 ng/μl of DNA (625-fold dilution). Challenging samples typically include fecal samples, which are contaminated with non-target DNA (*e.g.* plant and bacterial origin) and show elevated levels of overall DNA degradation. Therefore, we analyzed the performance of four DNA samples collected from Alpine ibex feces in the Gran Paradiso National Park. Three of the four fecal samples had a 25% genotyping rate (*i.e.* 248-270 genotyped loci). The fecal sample with the lowest quality had a 10.3% genotyping rate corresponding to 102 genotyped loci. Samples with low genotyping rates typically show inconsistently genotyped loci across samples. We found that this was indeed the case with 41-133 of the loci being genotyped in only one out of four fecal samples. A total of 41 loci were genotyped in at least three fecal samples (Figure 3D).

**Figure 3:**
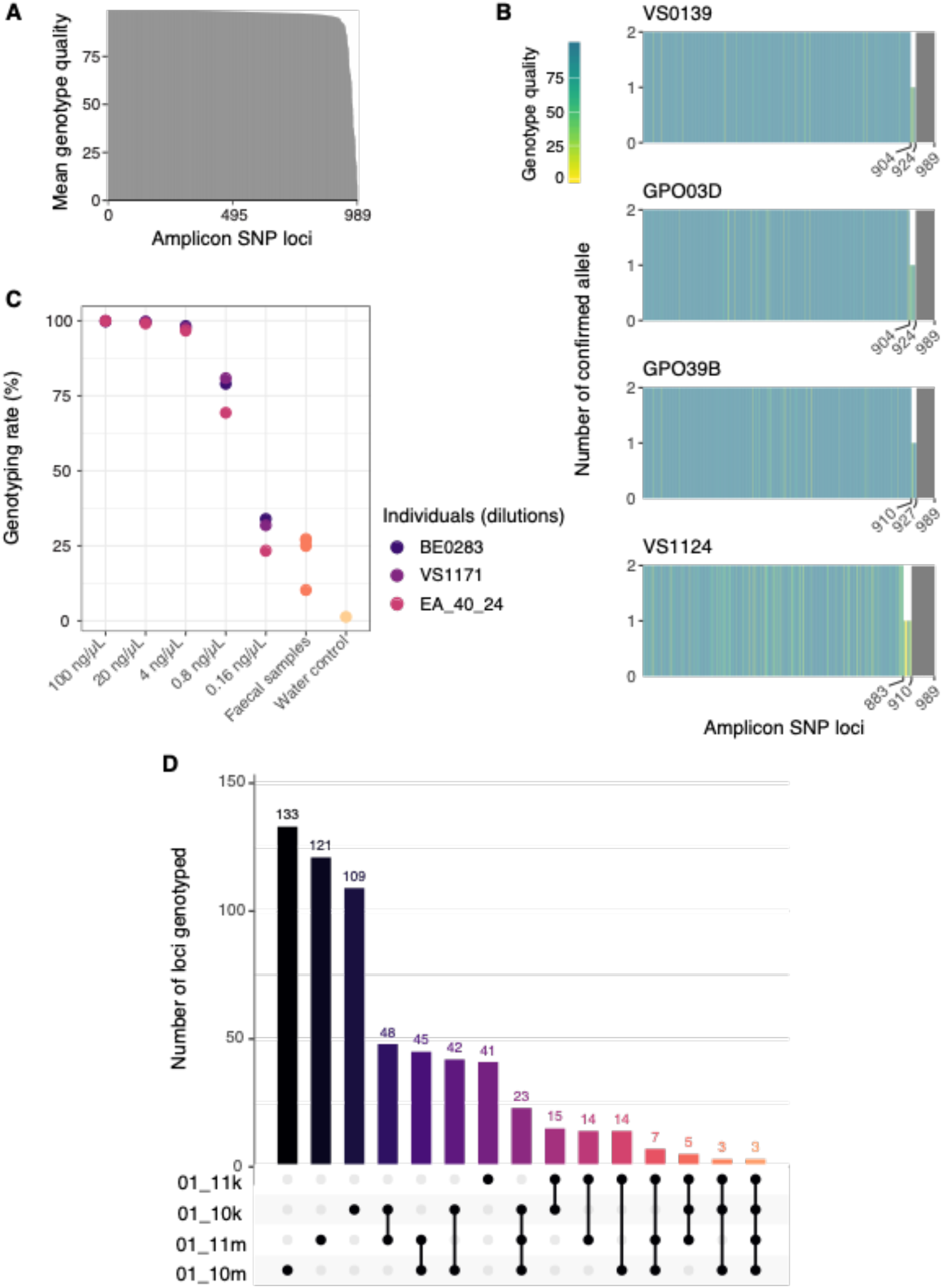
Genotyping accuracy and performance with low-quality or low-quantity input DNA. A) Mean genotyping quality (GQ) for each high-quality locus across the 65 high-quality DNA samples. B) Genotype accuracy assessed by matching recovered alleles from the targeted amplicon sequencing and whole-genome sequencing of the same four individuals. C) Assessment of genotyping rates for serial dilutions of three individuals and four fecal samples. D) Analysis of genotyped loci shared among the four fecal samples. The overlaps show recovered genotypes out of a total of 989 loci.

### High-resolution population structure

A major reason for genotyping species of conservation concern is the identification of population subdivisions and admixture events. We expanded our genotyping assay to the full set of Alpine ibex samples (n = 158) spanning the extant distribution range across the Alps. Based on a PCA, we identified three major genotype clusters. The largest cluster was composed of the Gran Paradiso source population and populations reintroduced directly from Gran Paradiso to Italy or Switzerland (Figure 4A). A second cluster grouped the two French populations Haute Maurienne and Champsaur (the latter founded with individuals coming from the former). The third cluster was composed of the isolated Alpi Marittime population, which is thought to have only six effective founder individuals (Terrier & Rossi, 1994). The identified population structure was also supported by individual ancestry coefficients using a sparse non-negative matrix factorization algorithm (K=3, Figure S1, Frichot et al. 2014). At the K with the lowest entropy (K=5, Figure S2), the analyses revealed a fine-scale population structure: all populations except for the two French populations Haute Maurienne and Champsaur, were clearly distinct (Figure 4C). The genotyping assay performed also well for population-level assignments of the low-input/quality samples. All four Gran Paradiso fecal samples had similar principal component values (Figure 4A) and structure population assignments (Figure 4C) as other, high quality samples from the same population. We also analyzed population differentiation of Alpine ibex using pairwise FST (Fig. 4B). We found a consistent pattern separating Alpi Marittime from all other populations. Furthermore, the two French populations showed relatively low differentiation consistent with their foundation history: the Champsaur population was founded only 25 years ago (less than 4 ibex generations) with 31 individuals coming from Haute Maurienne. The three Swiss populations (Albris, Brienzer Rothorn and Pleureur) were only weakly differentiated from the Gran Paradiso source population as expected from former analysis based on microsatellites (Biebach & Keller, 2009).

**Figure 4:**
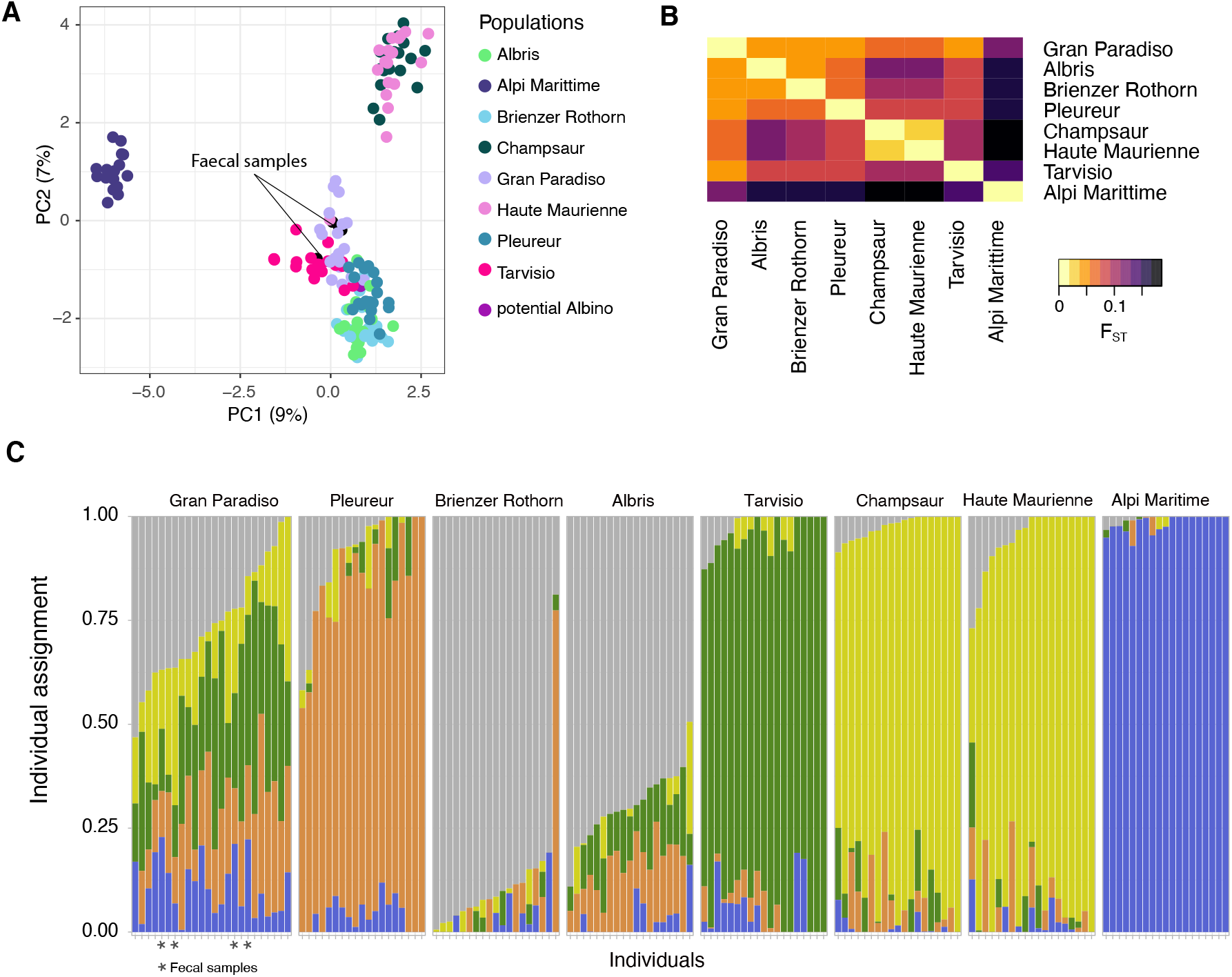
Population genetic analyses of Alpine ibex. A) Principal component analysis of 158 Alpine ibex individuals based on 617 genome-wide SNP loci designed for capturing genome-wide population differentiation. B) Pairwise F_ST_ matrix of all Alpine ibex populations based on genome-wide SNPs. Gran Paradiso was the only population surviving the near extinction and hence is the source population of all existing Alpine ibex populations. The Swiss populations Albris, Pleureur, Brienzer Rothorn and the Italian population Alpi Marittime were founded in early 1900 from Gran Paradiso individuals. The populations Tarvisio, Champsaur and Haute Maurienne were later founded independently from Gran Paradiso individuals. C) Structure-like analysis (based on sparse non-negative matrix factorization algorithms) of all Alpine ibex with K=5. *) fecal samples.

### Performance contrasts among next-generation sequencing methods

Next-generation sequencing methods for population monitoring can have distinct advantages or drawbacks. In contrast to the newly developed targeted amplicon sequencing for Alpine ibex, reduced representation (*e.g.* RAD-seq) and low-coverage whole genome sequencing have the potential for orders of magnitude larger numbers of scorable SNPs but of potentially lower genotyping quality and higher missingness. To objectively assess the performance of the newly developed assay, we analyzed genotyping outcomes of four core Alpine ibex populations (Gran Paradiso, Pleureur, Albris and Brienzer Rothorn; Figures 5, S2). After quality filtering of each dataset to ensure objective comparisons (see Methods), we retained 892 loci from targeted amplicon sequencing (*n* = 75 individuals), 26,547 RAD-seq loci (*n* = 82 individuals) and 3 Mio low-coverage whole genome sequencing loci (*n* = 15 individuals). Overall, 97% of the individuals had a per-individual genotyping rate of ≥90% for targeted amplicon sequencing loci contrasting with 23% of the individuals genotyped at ≥90 % for RAD-seq loci (Figure 5A). Enforcing a per-locus genotyping rate of ≥90% over all individuals, 96% of targeted amplicon sequencing loci but only 39% of RAD-seq loci were retained (Figure 5B). Locus and individual-level genotyping rates cannot meaningfully be retrieved from genotype likelihood-based analyses (low-coverage whole genome sequencing dataset). We performed comparative population differentiation analyses and found that the global F_ST_ ranged between 0.071 (RAD-seq) and 0.077 (targeted amplicon sequencing). Pairwise F_ST_ estimates were also similar among marker systems (Figure 5C). PCAs constructed from targeted amplicon sequencing and RAD-seq markers clearly resolved the four populations (Figure 5D, E). However, the low-coverage whole genome sequencing did not resolve Albris and Brienzer Rothorn populations (Figure 5F). The first and second principal component (PC) axes explained 6.2% and 4.9% for the targeted amplicon sequencing (Figure 5D), 5.2% and 4.2% for the RAD-seq (Figure 5E) and 23.8%-6.1% for the low-coverage whole genome sequencing (Figure 5F), respectively. Genotype assignments to clusters showed clear population differentiation for the targeted amplicon sequencing markers (Figure 5G) and slightly weaker resolution for RAD-seq markers (Figure 5H). Low-coverage whole genome sequencing genotyping clearly separated Gran Paradiso and Pleureur populations but again failed to resolve Albris and Brienzer Rothorn populations (Figure 5I).

**Figure 5:**
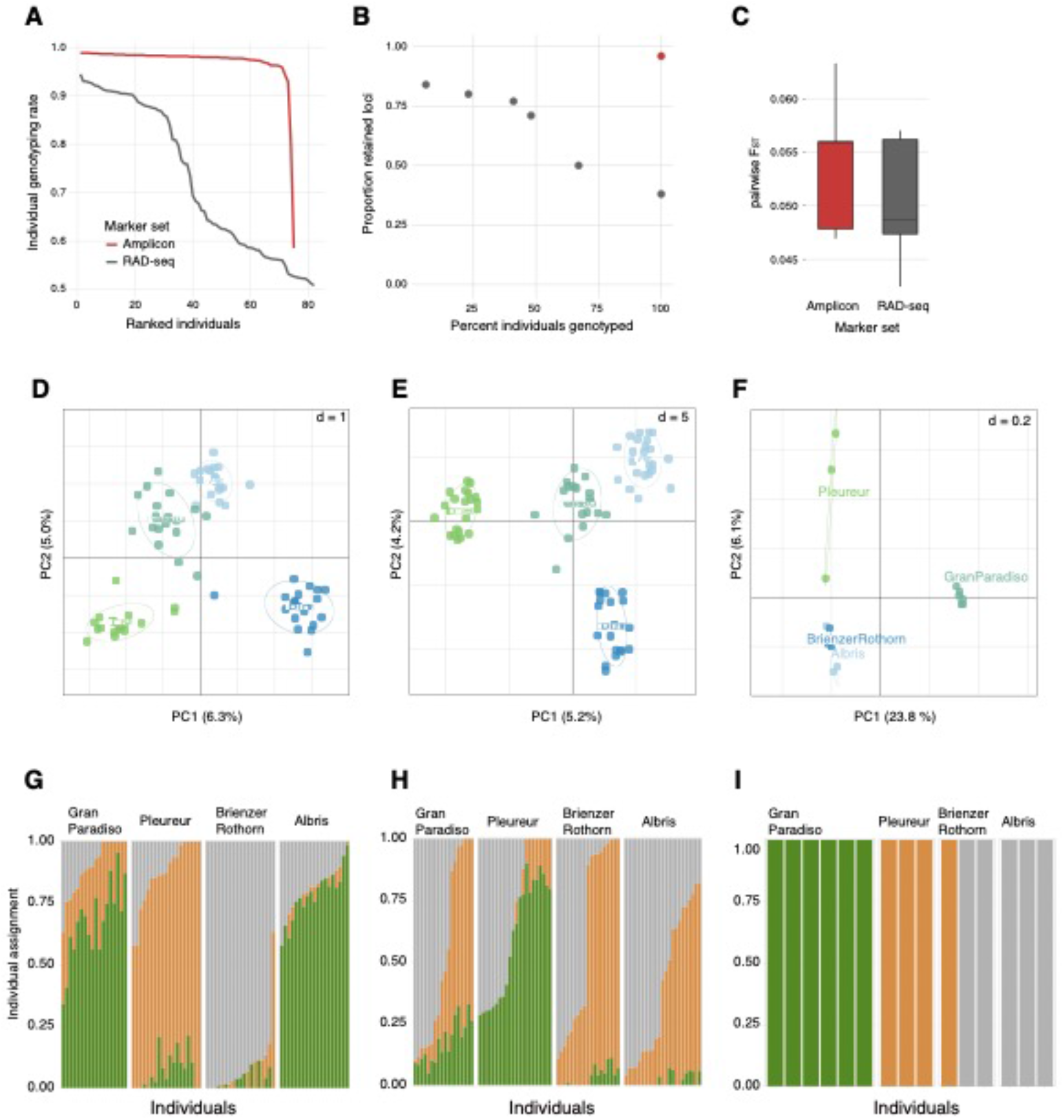
Performance comparisons between targeted amplicon sequencing, RAD-seq and low-coverage whole genome sequencing on four Alpine ibex populations. A) Individual genotyping rate ranked by individual for amplicon sequencing and RAD-seq. B) Proportion of SNP loci retained at a genotyping rate of 90% as a function of the percent of individuals included. The comparison is only meaningful for amplicon sequencing and RAD-seq. C) Boxplot of all pairwise FST estimates among populations for the amplicon sequencing and RAD-seq. D-F) Principal component analysis of population differentiation for (D) the amplicon sequencing (plotted are -PC1 against -PC2), (E) RAD-seq and (F) low-coverage whole-genome sequencing. Structure-like analysis (based on sparse non-negative matrix factorization algorithms) of (G) the amplicon sequencing, (H) RAD-seq and (I) low-coverage whole-genome sequencing datasets. I) is based on a NGSadmix analysis (Skotte et al., 2013).

### Detection of recent hybrids and introgression tracts

Introgression from domesticated animals is a major concern for a number of wild species. A set of SNP markers in the set of amplicons were specifically designed to detect introgression from domestic goats into Alpine ibex. To assess the power to discriminate genotypes suspected to be from hybrid individuals, we performed a PCA including all 158 Alpine ibex individuals, suspected hybrids based on field reports of unusual phenotypes (*n* = *6*), domestic goats (*n* = 5) and Iberian ibex (*n* = 3; Figure 6A). As expected, the first PC clearly separated domestic goats, Iberian ibex, as well as Alpine ibex. The second PC differentiated the Alpi Marittime population from all other Alpine ibex. The suspected hybrid individual from Alpi Marittime (AM_H) clearly clustered with domestic goats. Two additional suspected hybrids (GR_ib1, GR_ib2), as well as a potential albino individual (FR_blanc has largely whitish fur, but no red eyes) clustered with Alpine ibex. Three suspected hybrids (TI_ib, GPHB1 and GP_ib_V02_17) were located near the mid-point between domestic goats and Alpine ibex matching expectations for recent (F1 or backcross) hybrid genotypes.

**Figure 6:**
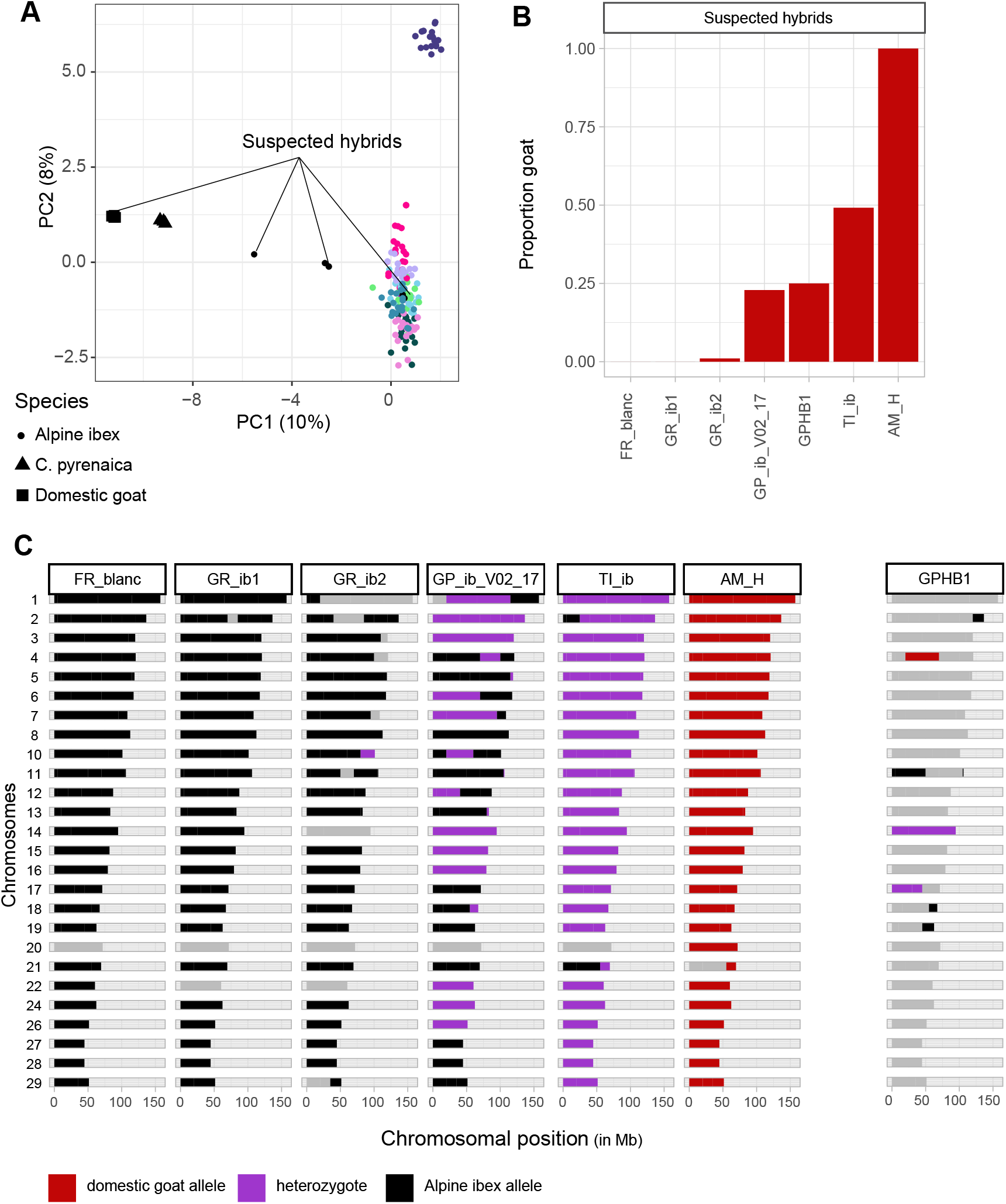
Performance of targeted amplicon sequencing to detect recent hybridization events. A) Principal component analysis of 168 individuals (excluding fecal samples) including domestic goat and Iberian ibex (N=5 and 3) and suspected hybrids based on phenotypic observations (N=6). The PCA was performed based on 617 genome-wide SNP loci designed for capturing putatively neutral population differentiation. B) Analysis of diagnostic markers for the detection of hybrids for 6 suspected hybrids and one potential albino. The proportion of diagnostic markers assigned to domestic goat are shown, where the proportion is calculated as (1*homozyogous goat genotype + ½*heterozygous genotype)/(total genotypes at diagnostic markers). C) Chromosome painting using diagnostic markers along the 29 autosomes. Colors identify Alpine ibex (blue), heterozygous (purple) and domestic goat (red) genotypes. Sample GPHB1 was of low DNA quality and quantity (drop of blood conserved on FTA filter paper; see Methods).

Using the goat-Alpine ibex diagnostic marker set, we analyzed the recency of the hybridization event by identifying contributions from each parental species (Figure 6B). Of the six suspected hybrids and one potential albino, two individuals were confirmed to be Alpine ibex (FR_blanc, GR_ib1) and one a domestic goat (AM_H). Our results confirm that the unusual phenotype of FR_blanc reported from the field was not caused by domestic goat introgression. Individual GR_ib1 was suspected to be a hybrid because it was behaving in very unusual ways, seeking proximity to buildings. Individual AM_H was reported to resemble domestic goat but was living among Alpine ibex. Another suspected hybrid (GR_ib2) with white hoofs showed a weak sign of domestic goat introgression (only one marker on chromosome 10 was heterozygous) suggesting a potential backcross. Note that GR_ib2 showed no clear differentiation from Alpine ibex based on the genome-wide marker PCA underlining the usefulness of specifically designed diagnostic loci. Two individuals (GP_ib_V02_17 and GPHB) showed signs of ~25% domestic goat introgression (*i.e.* likely F2 backcrosses). GP_ib_V02_17 had darker fur than commonly seen in Alpine ibex and an unusual horn shape (no nodes and a triangular transverse section). Chromosome painting showed one individual was being heterozygous for all diagnostic markers (TI_ib, Figure 6C), hence representing very likely an F1 hybrid. This individual was observed going into a stable following goats and had an unusual horn shape. The DNA from GPHB was from a small amount of blood stored on FTA paper explaining the poor genotyping quality. Field reports suspected a F1 hybrid but both the contribution plot (Figure 6B) and the chromosomal painting (Figure 6C) suggest a F2 hybrid. Our analyses show the power of highly discriminatory markers to detect recent hybridization and introgression events in a pool of individuals with field-reported, suspected admixture.

### Immunogenetics of Alpine ibex populations

The targeted amplicon sequencing specifically focused on 297 polymorphisms in immune-related genes within and outside the MHC. Using whole-genome sequencing data for Alpine ibex (*n* = 29), we found that the Tajima’s *D* in Alpine ibex ranged from −2.4 to 4.5 across all genes encoded in the MHC region and from −1.4 to 2.5 for all immune-related genes outside of the MHC region targeted by the amplicon sequencing. The median Tajima’s *D* was lower in the MHC (0.45) compared to immune loci outside of the MHC (1.3, Figure 7A). The high Tajima’s *D* values in some immune-related loci suggest long-term maintenance of alleles through balancing selection (Figure 7A). Comparing different sets of amplicon targets in our assay, we found that the genome-wide markers aimed at resolving population structure showed overall the highest average heterozygosity (Figure 7C). The lowest genome-wide heterozygosity was found in Alpi Marittime consistent with the severe founding bottleneck. Immune-related and MHC loci showed consistently lower levels of heterozygosity compared to genome-wide loci. With a notable exception of the Alpi Marittime population where the MHC showed higher levels of heterozygosity (0.25) than other immune-related and genome-wide markers (Figure 7C). The MHC of the Albris population showed a surprisingly low average heterozygosity (0.11) compared to genome-wide markers. On a PCA, the MHC in domestic goat and Iberian ibex showed a low degree of differentiation (Figure 7D). This is likely explained by the focus on segregating polymorphism in Alpine ibex only. MHC genotypes showed tight clusters among Alpine ibex individuals but only weak population signatures, which is in marked contrast to genome-wide markers (Figure 4A, 7D). Clusters of nearly identical Alpine ibex genotypes were generally composed of genotypes from multiple populations. The two French populations Haute Maurienne and Champsaur shared most MHC genotypes. Individuals from the Alpi Marittime were largely distinct from all other populations with the exception of a shared genotype with the Italian population Tarvisio. Some individuals from Alpi Marittime were translocated to Tarvisio in 1993, which may explain our finding.

**Figure 7:**
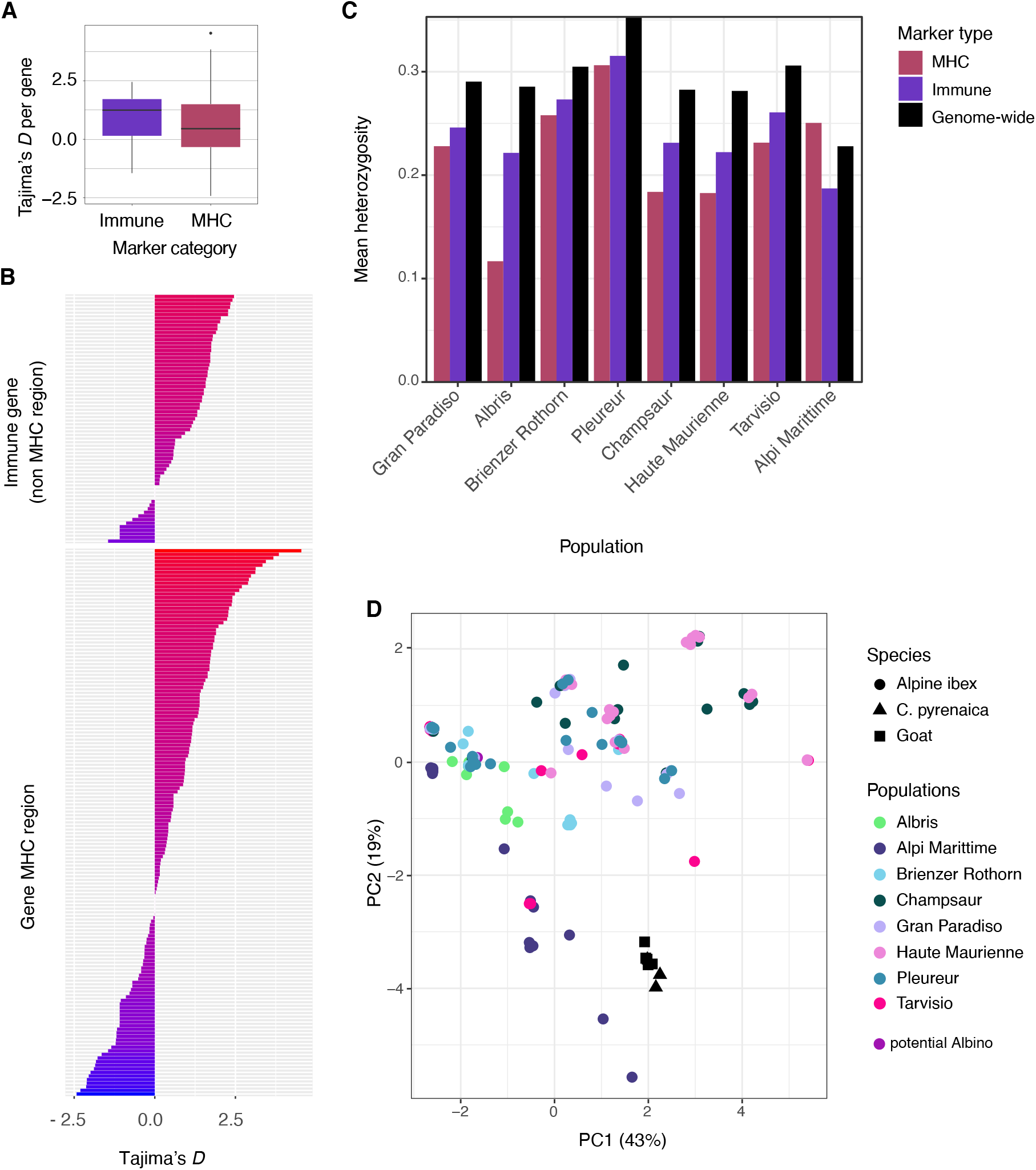
Genome-wide immunogenetics survey of Alpine ibex populations. A) Boxplot of Tajima’s *D* estimates based on whole-genome sequencing of 29 Alpine ibex of immune-related loci covered by the targeted amplicon sequencing assay. The MHC region is shown separately from other immune-related loci. B) Tajima’s *D* estimates shown per each gene represented by immune-related loci covered by the targeted amplicon sequencing assay. C) Targeted amplicon sequencing analyses of heterozygosity across Alpine ibex populations (*N*=*154*, excluding fecal samples). Heterozygosity is shown separately for genome-wide SNPs, the MHC region and other immune-related SNP loci. D) Principal component analysis of Alpine ibex and sister species based on 138 SNPs genotyped in the MHC region using targeted amplicon sequencing.

## Discussion

Genetic monitoring is central to many population surveys and conservation efforts. However, there is currently no established implementation enabling both targeted and highly parallelized genotyping. Here, we developed an accurate and versatile tool based on whole-genome sequence derived SNP loci for the monitoring of Alpine ibex population health. The set of nearly a thousand loci enables the concurrent assessment of population structure, the detection of recent hybridization events and immune function related genotypes. We show that our new amplicon array performs well even with low-input samples. The amplicon sequencing also performs at least equally well as other widely used population genomic approaches while providing key implementation advantages.

Effective high-throughput genotyping relies on the consistent amplification of a large number of loci and robustness to variation in input DNA quality. We show that the targeted sequencing approach produces largely uniform coverage both across a thousand loci and nearly 200 samples of highly variable DNA quantity and quality. Integrating whole-genome sequencing derived SNPs from different species into the design of amplicons allowed us to amplify across Alpine ibex, Iberian ibex and the domestic goat. This extends the usability of the assay with the caveat that the potential ascertainment bias in other species should be considered because SNPs were selected based on polymorphism in Alpine ibex only. Furthermore, the identified genotypes were well-validated by cross-referencing with the whole-genome sequencing datasets. Uniform, high-accuracy genotyping with low input DNA has previously only been achieved through microsatellite marker analyses (Hodel et al., 2016) or SNP chips designed on related model species (Cronin et al., 2015; Pertoldi et al., 2009). Compared to amplicon sequencing, SNP chips often lack extensibility and provide no sequence information surrounding the genotyped loci. An application of SNP chip genotyping in Alpine ibex by (Grossen et al., 2014) was based on a 52 K Illumina Goat SNP Chip (Tosser-Klopp et al., 2015). Among 95 Alpine ibex individuals, the study recovered 677 polymorphic markers out of a total of ~52’000 markers known to be polymorphic among domestic goats. This shows how even SNP chips designed for closely related species can be largely unsuitable and may suffer from substantial ascertainment bias. High-throughput methods such as RAD-seq and GBS sequencing are the most widely used next-generation sequencing approaches for non-model species often producing at least tens of thousands of SNPs. However, such restriction-based reduced representation approaches do not allow the targeting of specific loci (e.g. immune-relevant loci) and often produce highly uneven read depth across loci (Jiang et al., 2016). Furthermore, we show that replicating RAD-seq and low-coverage whole genome sequencing on the same set of populations analyzed in our targeted amplicon sequencing approach produces a highly similar resolution of the genetic structure. The comparatively low number of SNPs assayed in the targeted amplicon sequencing is most likely compensated by the highly consistent genotyping rates across loci and individuals.

We show that the microfluidics approach for amplicon sequencing produces consistent amplification across nearly all loci if minimum DNA requirements are fulfilled. We have set a minimum threshold of 50 ng/μl (or a total of 100 ng), which is usually achievable for well preserved tissue samples. We have also investigated the potential to recover genotypes from degraded and low-concentration samples such as fecal material. Fecal sampling is a widely used non-invasive sampling method and sometimes the only option for elusive species (Beja-Pereira et al., 2009). Fecal samples produce low quality (degradation) and low quantity DNA. In addition, fecal DNA samples can be heavily contaminated with bacteria, plant or prey DNA. Hence, the actual endogenous DNA is often much below the measured total DNA concentration. We analyzed fecal DNA quantities from 4-23 ng/μl but found that the proportion of successfully amplified loci was comparable to diluted non-fecal DNA at around 0.1 ng/μl. Consequently, non-targeted sequencing methods (*e.g.* RAD or whole-genome sequencing) would require very significant sequencing depth to adequately genotype endogenous DNA of such samples, because a large proportion of reads would be lost to non-endogenous DNA. We show that if analyses of fecal samples are combined with good quality samples, genotypes recovered from fecal DNA can be reliably assigned to individual populations. Furthermore, multiple amplifications of the same fecal samples could result in even better overall genotyping rates. The broad range of acceptable input DNA makes the targeted amplification of selected polymorphisms widely applicable across study systems provided that genomic resources from a small number of individuals are available for the assay development.

To assess the resolution of the genome-wide set of amplicon markers, we have genotyped a collection of Alpine ibex samples representing all major reintroduction events. Alpine ibex were limited to the Gran Paradiso National Park in Italy in the 19^th^ century. Starting in the early 20^th^ century, populations were introduced independently from Gran Paradiso to Switzerland, France and Italy and from there further populations were founded. The population structure of extant Alpine ibex populations are dominated by signals of population reintroductions and translocations (Biebach & Keller, 2009; Grossen et al., 2018). The Alpine ibex genotypes assessed by our targeted sequencing approach confirmed all major aspects of the reintroduction history including placing Gran Paradiso at the center of the extant genetic diversity. Populations reintroduced directly from Gran Paradiso were at the closest genetic distances to the source population yet showed distinct clustering as expected from the strong bottlenecks imposed by the translocation of few individuals. A major concern for wild species co-occurring with closely related domestic animals is the potential for hybridization and introgression. Alpine ibex populations across Europe are monitored for the presence of individuals with atypical phenotypes (e.g. Giacometti et al., 2004; Steyer et al., 2016; Todesco et al., 2016). We assessed genotypes of seven suspected hybrids (including one potential albino) and could show that only three individuals are clearly identifiable as recent hybrids. One individual showed possible signs of past introgression and three suspected hybrids clustered either with domestic goat or Alpine ibex genotypes highlighting the importance of genetic monitoring of suspected hybridization events.

A major genetic factor for the long-term survival of endangered species is diversity at important immune loci, in particular the MHC. We successfully amplified hundreds of loci involved in key immune functions focusing on non-synonymous polymorphisms. In parallel, we amplified a dense array of loci spanning the MHC. This 2-Mb locus on chromosome 23 recently received genetic material from domestic goats regenerating heterozygosity at the DRB locus (Grossen et al., 2014). Genotypes across the MHC clustered tightly among populations revealing that Alpine ibex were genetically impoverished across the entire MHC. Comparative analyses with domestic goat breeds will enable a refinement of our understanding of historic and potentially ongoing introgression into the Alpine ibex gene pool. Our findings of low diversity at the MHC are also in accordance with previous studies based on microsatellite and RAD genotyping (*e.g*. (e.g. Alasaad et al., 2012; Brambilla et al., 2018; Grossen et al., 2014). Underlining the relevance of performing large-scale targeted amplicon sequencing of immune loci are recent findings that high heterozygosity at the MHC was correlated with higher resistance to infectious keratoconjunctivitis, a major disease factor in some Alpine ibex populations (Brambilla et al., 2018). Surveillance of disease susceptibility through genetic analyses of immune loci will also significantly improve conservation strategies by informing the choice of founder individuals prior to reintroductions.

In conclusion, our highly parallel targeted amplicon assay demonstrates how next-generation sequencing techniques can be adapted for the needs of population genetic surveys and conservation monitoring. The WGS-informed approach enabled selecting a highly specific set of loci to simultaneously address questions of population structure, recent hybridization events and how polymorphism is shaped across major components of the immune system. Efficient and precise characterization of individual genotypes can be translated into recommendations on how to prioritize translocation events and replenish genetic diversity at immune loci. The versatility to amplify the same loci across related species enables also powerful screens for recent introgression events. Our study shows the relevance of bridging population genomic investigations with assays that can be realistically implemented into population genetic surveys and decision-making for conservation management.

## Acknowledgements

We are thankful to the scientific teams of the Gran Paradiso National Park, Parco Naturale Alpi Marittime, Parc National des Ecrins, Parc National de la Vanoise and Ente Foreste Tarvisio, for providing us with DNA samples from their Alpine ibex populations. Xenia Wietlisbach helped prepare samples. Targeted DNA sequencing and preparation was carried out at the Genetic Diversity Center (GDC) at ETHZ. This study makes use of data generated by the NextGen Consortium, which was supported by grant agreement number 244356 of the European Union’s Seventh Framework Programme (FP7/2010-2014). This study was performed in the framework of the EU funded Interreg Alcotra V-A France-Italy 1664 LEMED-IBEX.

## Data accessibility

The raw amplicon sequencing data produced for this project was deposited at the NSBI Short Read Archive under the Bioproject Accession number PRJNA669599. The raw whole-genome sequencing data produced for this project was deposited at the NCBI Short Read Archive under the Accession nos. SAMN10736122–SAMN10736160 (BioProject PRJNA514886 [https://www.ncbi.nlm.nih.gov/sra/PRJNA514886]). The whole-genome data produced by the NexGen Consortium (Capra hircus accessions: ERR470105, ERR470101, ERR313212, ERR313211, ERR313204, ERR297229, ERR313206, ERR405774, ERR405778, ERR315778, ERR318768, ERR246140, ERR340429, ERR246152, ERR345976, Capra aegagrus accessions: ERR340334, ERR340340, ERR340333, ERR340331, ERR340335, ERR340348; Ovis aries accessions: ERR157945, ERR299288; Ovis orientalis: ERR157938; Ovis vignei: ERR454948; Ovis canadensis: SRR501898) was downloaded from [ftp://ftp.sra.ebi.ac.uk/vol1/fastq].

## Author contributions

CK, CG and DC conceived the study, CK, DW, GC, IB and CG performed analyses, AB contributed samples, DL contributed datasets, CK, CG and DC wrote the manuscript

## Supplementary Information

(SupplementaryTables.xlsx)

**Supplementary Table S1: Overview of individual sampling.** Overview of all sequenced individuals analysed in this study including information of species, population, inclusion in high-quality samples and sample type.

**Supplementary Table S2: Overview of SNP positions included in the marker design.** Shown are marker type (Genome-wide, Diagnostic, MHC region, Immune), Batch (1st or 2nd submission to Fluidigm for marker design), Exclusion (N: Kept in final marker set, Y: Yes, excluded due to quality checks), Excluded_by (N_percentage: contained >50% masked positions in the sequence surrounding the focal SNP; Fluidigm: marker did not pass Fluidigm design principles; VariantFiltration: marker failed GATK hard filtering; Non_variant: marker was invariant; <75%: genotyping rate was below 75% among individuals), functional annotation is based on SnpEff.

**Supplementary Table S3: Individual read numbers**. Given are read numbers per each individual before and after quality trimming, merging and genome alignment.

**Supplementary Figure 1:**
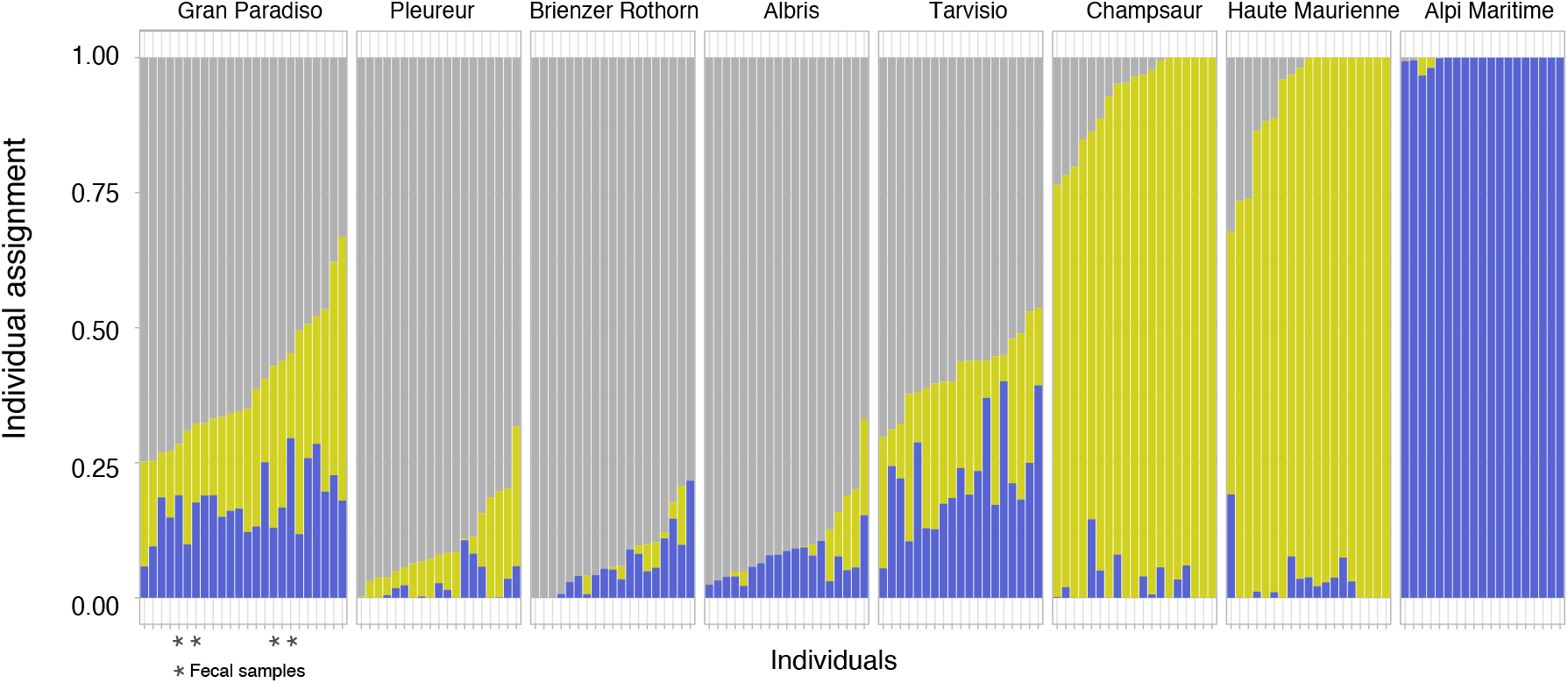
Population genetic analyses of Alpine ibex. Structure-like visualization (based on sparse non-negative matrix factorization algorithms) of all Alpine ibex with *K*=3.

**Supplementary Figure 2:**
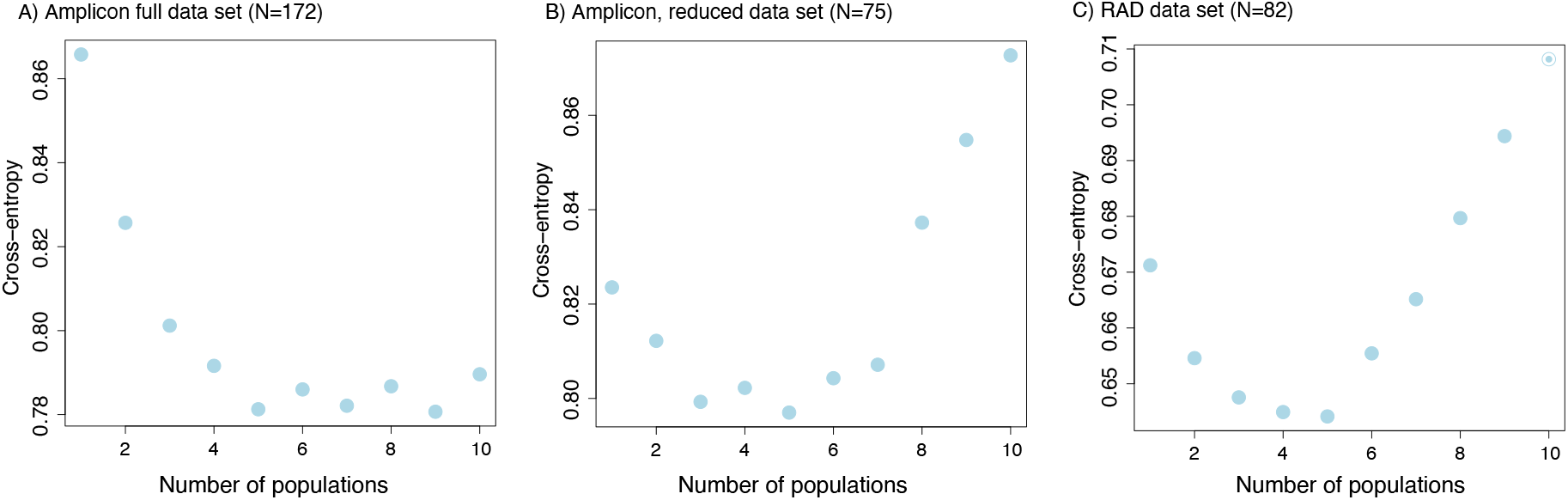
Structure-like analysis of Alpine ibex. Shown is the distribution of entropy for K varied from 1 to 10 (100 replicates each) for the Structure-like analysis (sparse non-negative matrix factorization algorithms) based on (A) the full Alpine ibex amplicon data set, (B) the reduced amplicon data (for methods comparison) and (C) the RAD data.

## References

Acevedo-Whitehouse, K., & Cunningham, A. A. (2006). Is MHC enough for understanding wildlife immunogenetics? Trends in Ecology & Evolution, 21(8), 433–438.

Alasaad, S., Biebach, I., Grossen, C., Soriguer, R. C., Pérez, J. M., & Keller, L. F. (2012). Microsatellite-based genotyping of MHC class II DRB1 gene in Iberian and Alpine ibex. European Journal of Wildlife Research, 58(4), 743–748.

Allendorf, F. W., Hohenlohe, P. A., & Luikart, G. (2010). Genomics and the future of conservation genetics. Nature Reviews. Genetics, 11(10), 697–709.

Altschul, S. F., Gish, W., Miller, W., Myers, E. W., & Lipman, D. J. (1990). Basic local alignment search tool. Journal of Molecular Biology, 215(3), 403–410.

Ammerdorffer, A., Roest, H.-I. J., Dinkla, A., Post, J., Schoffelen, T., van Deuren, M., Sprong, T., & Rebel, J. M. (2014). The effect of C. burnetii infection on the cytokine response of PBMCs from pregnant goats. PloS One, 9(10), e109283.

Angelone, S., Jowers, M. J., Molinar Min, A. R., Fandos, P., Prieto, P., Pasquetti, M., Cano-Manuel, F. J., Mentaberre, G., Olvera, J. R. L., Ráez-Bravo, A., Espinosa, J., Pérez, J. M., Soriguer, R. C., Rossi, L., & Granados, J. E. (2018). Hidden MHC genetic diversity in the Iberian ibex (Capra pyrenaica). BMC Genetics, 19(1), 28.

Barnosky, A. D., Matzke, N., Tomiya, S., Wogan, G. O. U., Swartz, B., Quental, T. B., Marshall, C., McGuire, J. L., Lindsey, E. L., Maguire, K. C., Mersey, B., & Ferrer, E. A. (2011). Has the Earth’s sixth mass extinction already arrived? Nature, 471(7336), 51–57.

Beja-Pereira, A., Oliveira, R., Alves, P. C., Schwartz, M. K., & Luikart, G. (2009). Advancing ecological understandings through technological transformations in noninvasive genetics. Molecular Ecology Resources, 9(5), 1279–1301.

Bernatchez, L., & Landry, C. (2003). MHC studies in nonmodel vertebrates: what have we learned about natural selection in 15 years? Journal of Evolutionary Biology, 16(3), 363–377.

Bickhart, D. M., Rosen, B. D., Koren, S., Sayre, B. L., Hastie, A. R., Chan, S., Lee, J., Lam, E. T., Liachko, I., Sullivan, S. T., Burton, J. N., Huson, H. J., Nystrom, J. C., Kelley, C. M., Hutchison, J. L., Zhou, Y., Sun, J., Crisà, A., Ponce de León, F. A., … Smith, T. P. L. (2017). Single-molecule sequencing and chromatin conformation capture enable de novo reference assembly of the domestic goat genome. Nature Genetics, 49(4), 643–650.

Biebach, I., & Keller, L. F. (2009). A strong genetic footprint of the re-introduction history of Alpine ibex (Capra ibex ibex). Molecular Ecology, 18(24), 5046–5058.

Bolger, A. M., Lohse, M., & Usadel, B. (2014). Trimmomatic: a flexible trimmer for Illumina sequence data. Bioinformatics, 30(15), 2114–2120.

Bollmer, J. L., Vargas, F. H., & Parker, P. G. (2007). Low MHC variation in the endangered Galápagos penguin (Spheniscus mendiculus). Immunogenetics, 59(7), 593–602.

Brambilla, A., Biebach, I., Bassano, B., Bogliani, G., & von Hardenberg, A. (2015). Direct and indirect causal effects of heterozygosity on fitness-related traits in Alpine ibex. Proceedings. Biological Sciences / The Royal Society, 282(1798), 20141873.

Brambilla, A., Keller, L., Bassano, B., & Grossen, C. (2018). Heterozygosity-fitness correlation at the major histocompatibility complex despite low variation in Alpine ibex (*Capra ibex*). Evolutionary Applications, 11(5), 631–644.

Brambilla, A., Von Hardenberg, A., Nelli, L., & Bassano, B. (2020). Distribution, status, and recent population dynamics of Alpine ibex Capra ibex in Europe. Mammal Review, 50(3), 267–277.

Ceballos, G., Ehrlich, P. R., Barnosky, A. D., Garcia, A., Pringle, R. M., & Palmer, T. M. (2015). Accelerated modern human-induced species losses: Entering the sixth mass extinction. Science Advances, 1(5), e1400253–e1400253.

Ceballos, G., Ehrlich, P. R., & Dirzo, R. (2017). Biological annihilation via the ongoing sixth mass extinction signaled by vertebrate population losses and declines. Proceedings of the National Academy of Sciences, 201704949.

Cingolani, P., Platts, A., Wang, L. L., Coon, M., Nguyen, T., Wang, L., Land, S. J., Lu, X., & Ruden, D. M. (2012). A program for annotating and predicting the effects of single nucleotide polymorphisms, SnpEff: SNPs in the genome of Drosophila melanogaster strain w1118; iso-2; iso-3. Fly, 6(2), 80–92.

Cronin, M. A., Cánovas, A., Bannasch, D. L., Oberbauer, A. M., & Medrano, J. F. (2015). Single Nucleotide Polymorphism (SNP) Variation of Wolves (Canis lupus) in Southeast Alaska and Comparison with Wolves, Dogs, and Coyotes in North America. The Journal of Heredity, 106(1), 26–36.

Davey, J. W., Hohenlohe, P. A., Etter, P. D., Boone, J. Q., Catchen, J. M., & Blaxter, M. L. (2011). Genome-wide genetic marker discovery and genotyping using next-generation sequencing. Nature Reviews. Genetics, 12(7), 499–510.

Dong, Y., Xie, M., Jiang, Y., Xiao, N., Du, X., Zhang, W., Tosser-Klopp, G., Wang, J., Yang, S., Liang, J., Chen, W., Chen, J., Zeng, P., Hou, Y., Bian, C., Pan, S., Li, Y., Liu, X., Wang, W., … Wang, W. (2013). Sequencing and automated whole-genome optical mapping of the genome of a domestic goat (Capra hircus). Nature Biotechnology, 31(2), 135–141.

Ewels, P., Magnusson, M., Lundin, S., & Käller, M. (2016). MultiQC: summarize analysis results for multiple tools and samples in a single report. Bioinformatics, 32(19), 3047–3048.

Frankham, R. (2005). Genetics and extinction. Biological Conservation, 126(2), 131–140.

Garvin, M. R., Saitoh, K., & Gharrett, A. J. (2010). Application of single nucleotide polymorphisms to non-model species: a technical review. Molecular Ecology Resources, 10(6), 915–934.

Giacometti, M., Roganti, R., Tann, D. D., Stahlberger-Saitbekova, N., & Obexer-Ruff, G. (2004). Alpine ibex *Capra ibex ibex* x domestic goat *C. aegagrus domestica* hybrids in a restricted area of southern Switzerland. Wildlife Biology, 10(1), 137–143.

Grodinsky, C., & Stüwe, M. (1987). With lots of help alpine ibex return to their mountains. Smithsonian, 18(9), 68–77.

Grossen, C., Biebach, I., Angelone-Alasaad, S., Keller, L. F., & Croll, D. (2018). Population genomics analyses of European ibex species show lower diversity and higher inbreeding in reintroduced populations. Evolutionary Applications, 11(2), 123–139.

Grossen, C., Guillaume, F., Keller, L. F., & Croll, D. (2020). Purging of highly deleterious mutations through severe bottlenecks in Alpine ibex. Nature Communications. https://www.nature.com/articles/s41467-020-14803-1

Grossen, C., Keller, L. F., Biebach, I., International Goat Genome Consortium, & Croll, D. (2014). Introgression from domestic goat generated variation at the major histocompatibility complex of Alpine ibex. PLoS Genetics, 10(6), e1004438.

Hodel, R. G. J., Segovia-Salcedo, M. C., Landis, J. B., Crowl, A. A., Sun, M., Liu, X., Gitzendanner, M. A., Douglas, N. A., Germain-Aubrey, C. C., Chen, S., Soltis, D. E., & Soltis, P. S. (2016). The Report of My Death was an Exaggeration: A Review for Researchers Using Microsatellites in the 21st Century. Applications in Plant Sciences, 4(6), 1600025.

Holderegger, R., Balkenhol, N., Bolliger, J., Engler, J. O., Gugerli, F., Hochkirch, A., Nowak, C., Segelbacher, G., Widmer, A., & Zachos, F. E. (2019). Conservation genetics: Linking science with practice. Molecular Ecology, 28(17), 3848–3856.

Jiang, Z., Wang, H., Michal, J. J., Zhou, X., Liu, B., Woods, L. C. S., & Fuchs, R. A. (2016). Genome Wide Sampling Sequencing for SNP Genotyping: Methods, Challenges and Future Development. International Journal of Biological Sciences, 12(1), 100–108.

Kim, D., Paggi, J. M., Park, C., Bennett, C., & Salzberg, S. L. (2019). Graph-based genome alignment and genotyping with HISAT2 and HISAT-genotype. Nature Biotechnology, 37(8), 907–915.

Korneliussen, T. S., Albrechtsen, A., & Nielsen, R. (2014). ANGSD: Analysis of Next Generation Sequencing Data. BMC Bioinformatics, 15(1), 356.

Kosch, T. A., Silva, C. N. S., & Brannelly, L. A. (2019). Genetic potential for disease resistance in critically endangered amphibians decimated by chytridiomycosis. Animal: An International Journal of Animal Bioscience. https://zslpublications.onlinelibrary.wiley.com/doi/abs/10.1111/acv.12459

Langmead, B., & Salzberg, S. L. (2012). Fast gapped-read alignment with Bowtie 2. Nature Methods, 9(4), 357–359.

Leigh, D. M., Lischer, H. E. L., Grossen, C., & Keller, L. F. (2018). Batch effects in a multiyear sequencing study: False biological trends due to changes in read lengths. Molecular Ecology Resources, 18(4), 778–788.

Li, H., Handsaker, B., Wysoker, A., Fennell, T., Ruan, J., Homer, N., Marth, G., Abecasis, G., Durbin, R., & 1000 Genome Project Data Processing Subgroup. (2009). The Sequence Alignment/Map format and SAMtools. Bioinformatics, 25(16), 2078–2079.

Magoč, T., & Salzberg, S. L. (2011). FLASH: fast length adjustment of short reads to improve genome assemblies. Bioinformatics, 27(21), 2957–2963.

McKenna, A., Hanna, M., Banks, E., Sivachenko, A., Cibulskis, K., Kernytsky, A., Garimella, K., Altshuler, D., Gabriel, S., Daly, M., & DePristo, M. A. (2010). The Genome Analysis Toolkit: A MapReduce framework for analyzing next-generation DNA sequencing data. Genome Research, 20(9), 1297–1303.

Meisner, J., & Albrechtsen, A. (n.d.). Inferring Population Structure and Admixture Proportions in Low-Depth NGS Data. 13.

Mick, V., Le Carrou, G., Corde, Y., Game, Y., Jay, M., & Garin-Bastuji, B. (2014). Brucella melitensis in France: persistence in wildlife and probable spillover from Alpine ibex to domestic animals. PloS One, 9(4), e94168.

O’Brien, S. J., Johnson, W. E., Driscoll, C. A., Dobrynin, P., & Marker, L. (2017). Conservation Genetics of the Cheetah: Lessons Learned and New Opportunities. The Journal of Heredity, 108(6), 671–677.

O’Brien, S. J., Roelke, M., Marker, L., Newman, A., Winkler, C., Meltzer, D., Colly, L., Evermann, J., Bush, M., & Wildt, D. (1985). Genetic basis for species vulnerability in the cheetah. Science, 227(4693), 1428–1434.

O’Brien, S. J., Wildt, D. E., Goldman, D., Merril, C. R., & Bush, M. (1983). The Cheetah Is Depauperate in Genetic Variation. Science, 221(4609), 459–462.

Ouborg, N. J., Pertoldi, C., Loeschcke, V., Bijlsma, R. K., & Hedrick, P. W. (2010). Conservation genetics in transition to conservation genomics. Trends in Genetics: TIG, 26(4), 177–187.

Pertoldi, C., Tokarska, M., Wójcik, J. M., Demontis, D., Loeschcke, V., Gregersen, V. R., Coltman, D., Wilson, G. A., Randi, E., Hansen, M. M., & Bendixen, C. (2009). Depauperate genetic variability detected in the American and European bison using genomic techniques. Biology Direct, 4, 48; discussion 48.

Pimm, S. L., Dollar, L., & Bass, O. L. (2006). The genetic rescue of the Florida panther. Animal Conservation, 9(2), 115–122.

Quéméré, E., Rossi, S., Petit, E., Marchand, P., Merlet, J., Game, Y., Galan, M., & Gilot-Fromont, E. (2020). Genetic epidemiology of the Alpine ibex reservoir of persistent and virulent brucellosis outbreak. Scientific Reports, 10(1), 4400.

Quinlan, A. R., & Hall, I. M. (2010). BEDTools: a flexible suite of utilities for comparing genomic features. Bioinformatics, 26(6), 841–842.

Reed, D. H., & Frankham, R. (2003). Correlation between Fitness and Genetic Diversity. Conservation Biology: The Journal of the Society for Conservation Biology, 17(1), 230–237.

Roelke, M. E., Martenson, J. S., & O’Brien, S. J. (1993). The consequences of demographic reduction and genetic depletion in the endangered Florida panther. Current Biology: CB, 3(6), 340–350.

Savage, A. E., Sredl, M. J., & Zamudio, K. R. (2011). Disease dynamics vary spatially and temporally in a North American amphibian. Biological Conservation, 144(6), 1910–1915.

Schoville, S. D., Bonin, A., François, O., Lobreaux, S., Melodelima, C., & Manel, S. (2012). Adaptive Genetic Variation on the Landscape: Methods and Cases. Annual Review of Ecology, Evolution, and Systematics, 43(1), 23–43.

Siddle, H. V., Kreiss, A., Eldridge, M. D. B., Noonan, E., Clarke, C. J., Pyecroft, S., Woods, G. M., & Belov, K. (2007). Transmission of a fatal clonal tumor by biting occurs due to depleted MHC diversity in a threatened carnivorous marsupial. Proceedings of the National Academy of Sciences of the United States of America, 104(41), 16221–16226.

Skotte, L., Korneliussen, T. S., & Albrechtsen, A. (2013). Estimating Individual Admixture Proportions from Next Generation Sequencing Data. Genetics, 195(3), 693–702.

Steiner, C. C., Putnam, A. S., Hoeck, P. E. A., & Ryder, O. A. (2013). Conservation genomics of threatened animal species. Annual Review of Animal Biosciences, 1, 261–281.

Steyer, K., Kraus, R. H. S., Mölich, T., Anders, O., Cocchiararo, B., Frosch, C., Geib, A., Götz, M., Herrmann, M., Hupe, K., Kohnen, A., Krüger, M., Müller, F., Pir, J. B., Reiners, T. E., Roch, S., Schade, U., Schiefenhövel, P., Siemund, M., … Nowak, C. (2016). Large-scale genetic census of an elusive carnivore, the European wildcat (Felis s. silvestris). Conservation Genetics, 17(5), 1183–1199.

Tarasov, A., Vilella, A. J., Cuppen, E., Nijman, I. J., & Prins, P. (2015). Sambamba: fast processing of NGS alignment formats. Bioinformatics, 31(12), 2032–2034.

Terrier, G., & Rossi, P. (1994). Le bouquetin (Capra ibex ibex) dans les Alpes Maritimes Franco-Italiennes: occupation de l’espace, colonisation et régulation naturelles. Travaux Scientifiques Du Parc National de La Vanoise, 18, 271–287.

Todesco, M., Pascual, M. A., Owens, G. L., Ostevik, K. L., Moyers, B. T., Hübner, S., Heredia, S. M., Hahn, M. A., Caseys, C., Bock, D. G., & Rieseberg, L. H. (2016). Hybridization and extinction. Evolutionary Applications, 9(7), 892–908.

Tosser-Klopp, G., Rothschild, M. F., Huson, H. J., Nicolazzi, E. L., Sonstegard, T. S., Amills, M., Riggs, P., Van Tassell, C. P., Marsan, P. A., Stella, A., & Others. (2015). Update on the International Goat Genome Consortium Projects. Plant and Animal Genome. https://hal.inrae.fr/hal-02800945

Tschirren, B., Andersson, M., Scherman, K., Westerdahl, H., Mittl, P. R. E., & Råberg, L. (2013). Polymorphisms at the innate immune receptor TLR2 are associated with Borrelia infection in a wild rodent population. Proceedings. Biological Sciences / The Royal Society, 280(1759), 20130364.

Turner, A. K., Begon, M., Jackson, J. A., Bradley, J. E., & Paterson, S. (2011). Genetic Diversity in Cytokines Associated with Immune Variation and Resistance to Multiple Pathogens in a Natural Rodent Population. PLoS Genetics, 7(10), e1002343.

Van der Auwera, G. A., Carneiro, M. O., Hartl, C., Poplin, R., Del Angel, G., Levy-Moonshine, A., Jordan, T., Shakir, K., Roazen, D., Thibault, J., Banks, E., Garimella, K. V., Altshuler, D., Gabriel, S., & DePristo, M. A. (2013). From FastQ data to high confidence variant calls: the Genome Analysis Toolkit best practices pipeline. Current Protocols in Bioinformatics / Editoral Board, Andreas D. Baxevanis … [et Al.], 43, 11.10.1-11.10.33.

Willisch, C. S., Biebach, I., Koller, U., Bucher, T., Marreros, N., Ryser-Degiorgis, M.-P., Keller, L. F., & Neuhaus, P. (2012). Male reproductive pattern in a polygynous ungulate with a slow life-history: the role of age, social status and alternative mating tactics. Evolutionary Ecology, 26(1), 187–206.

Witzenberger, K. A., & Hochkirch, A. (2014). The genetic integrity of the ex situ population of the European wildcat (Felis silvestris silvestris) is seriously threatened by introgression from domestic cats (Felis silvestris catus). PloS One, 9(8), e106083.

Wwf. (2018). Living Planet Report - 2018: Aiming Higher (Grooten, M. and Almond, R.E.A.(Eds)). WWF.

Yates, A. D., Achuthan, P., Akanni, W., Allen, J., Allen, J., Alvarez-Jarreta, J., Amode, M. R., Armean, I. M., Azov, A. G., Bennett, R., Bhai, J., Billis, K., Boddu, S., Marugán, J. C., Cummins, C., Davidson, C., Dodiya, K., Fatima, R., Gall, A., … Flicek, P. (2019). Ensembl 2020. Nucleic Acids Research, gkz966.

Zhu, Y., Grueber, C., Li, Y., He, M., Hu, L., He, K., Liu, H., Zhang, H., & Wu, H. (2020). MHC - associated Baylisascaris schroederi load informs the giant panda reintroduction program. International Journal for Parasitology. Parasites and Wildlife, 12, 113–120.

Camille Kessler, Alice Brambilla, Dominique Waldvogel, Glauco Camenisch, Iris Biebach, Deborah M Leigh, Christine Grossen, Daniel Croll; 2020; Amplicon sequencing of Alpine ibex; NCBI SRA; BioProject PRJNA669599

Christine Grossen, Frederic Guillaume, Lukas F. Keller, Daniel Croll; 2020; Whole-genome sequencing of ibex species; NCBI SRA; BioProject PRJNA514886

NextGen consortium; Next generation methods to preserve farm animal biodiversity NEXTGEN; 2014; NCBI SRA; ERR470105, ERR470101, ERR313212, ERR313211, ERR313204, ERR297229, ERR313206, ERR405774, ERR405778, ERR315778, ERR318768, ERR246140, ERR340429, ERR246152, ERR345976, ERR340334, ERR340340, ERR340333, ERR340331, ERR340335, ERR340348, ERR157945, ERR299288, ERR157938, ERR454948, SRR501898

